# Alternative mRNA splicing controls the functions of the histone H3K27 demethylase UTX/KDM6A

**DOI:** 10.1101/2023.04.20.536794

**Authors:** Omid Fotouhi, Sheikh Nizamuddin, Stephanie Falk, Oliver Schilling, Ruth Knüchel-Clarke, Martin L. Biniossek, H.Th. Marc Timmers

**Affiliations:** Department of Urology, Medical Center-University of Freiburg, 79016 Freiburg, Germany; German Cancer Consortium (DKTK) partner site Freiburg, German Cancer Research Center (DKFZ), 69120 Heidelberg, Germany; Max Planck Institute of Immunobiology and Epigenetics, Freiburg, Germany; Institute for Surgical Pathology, Faculty of Medicine, Medical Center - University of Freiburg, University of Freiburg, Freiburg, Germany; Institute of Pathology, University Hospital RWTH Aachen, 52074 Aachen, Germany; Institute of Molecular Medicine and Cell Research, Faculty of Medicine, University of Freiburg, Freiburg, Germany

**Keywords:** alternative splicing, histone demethylase, histone methylation, cancer biology, chromatin, proteomics, bladder cancer

## Abstract

**Background:** The *UTX/KDM6A* histone H3K27 demethylase plays an important role in development and is frequently mutated in cancers such as urothelial cancer. Despite many studies on UTX proteins, variations in mRNA splicing have been overlooked.

**Methods:** Using Nanopore sequencing, we present a comprehensive analysis of *UTX/KDM6A* splicing events in human cell lines and in tissue samples from bladder cancer and normal epithelium.

**Results:** The central region of *UTX* mRNAs encoded by exons 12 to 17 undergoes extensive alternative splicing. Up to half of all stable mRNAs (8-48% in bladder tissues and 18-58% in cell lines) are represented by the *UTX* canonical isoform lacking exon 14 encoding a nuclear localization sequence, hence localize to the nucleus, unlike cytonuclear localization of the canonical isoform. Chromatin association was also higher for exon 14-containing isoform compared to the canonical UTX. Using quantitative mass spectrometry, we found that all UTX isoforms integrated into the MLL3 and MLL4, PR-DUB and MiDAC complexes. Interestingly, one of the novel UTX isoforms, which lacks exons 14 and 16, fails to interact with PR-DUB and MiDAC complex members.

**Conclusion:** *UTX* mRNAs undergo extensive alternative splicing that controls the subcellular localization of UTX and its interactions with other chromatin regulatory complexes.

**Simple Summary:** *UTX/KDM6A* is a histone H3K27 demethylase and plays an important role in mammalian development and human diseases such as urothelial cancer. We identified a region encompassing exons 12-17 of *UTX* that undergoes extensive splicing events. As a result, a nuclear localization sequence located in exon14 is missing in a considerable part of *UTX* transcripts in different cell lines and tissues from normal bladder epithelium and bladder cancer. Mass spectrometry analysis showed a role for this region in binding to the epigenetic PR-DUB and MiDAC complexes. UTX was also more extensively bound to chromatin when the alternative splicing region presented. Our study showed that alternative splicing of *UTX* transcripts plays an important role in its functions.

## Introduction

Chromatin modifiers play a critical role during normal differentiation and in disease states. The Ubiquitously Transcribed tetratricopeptide repeat (TPR) on chromosome X gene, *UTX* (also often referred to as *KDM6A*) encodes a histone H3K27me3 demethylase. Throughout the remainder of this manuscript, we will refer to *UTX/KDM6A* as *UTX* for simplicity. The *UTX* gene is highly mutated in different cancer types [1] and especially in bladder cancer (BLCA) with mutation frequencies of up to 30% [2,3]. Even normal bladder epithelium contains many discrete regions of clonally expanded cells that harbor independent mutations in *UTX* [4]. Such “morphologically normal” predisposition aberrations were not observed for other frequently mutated genes in BLCA such as *TP53* and *RB1*, which indicates that *UTX* inactivation is an early event in bladder carcinogenesis [5].

The pathophysiological significance of UTX is historically attributed to its C-terminal JmjC domain, which carries the histone demethylation function. However, UTX comprises two other functional regions, an N-terminal TPR region and an intrinsically disordered region (IDR) in the middle of the protein. The TPR is a highly conserved 34-40 amino acid motif tandem repeat, which is often involved in protein-protein interactions [6]. The IDR has been shown to be involved in the formation of biomolecular condensates [7]. UTX is an integral member of the MLL3 and MLL4 histone H3K4 methylation complexes of the COMPASS family of SET1/MLL complexes [8–10]. The N-terminal PHD region of MLL3/KMT2C has been shown to bind the BAP1-containing PR-DUB complex, which is also a tumor suppressor and acts as a deubiquitinase for histone H2A. PR-DUB binding is compromised by cancer mutations in MLL3 and could be partially balanced by UTX in an experimental setting [11]. Recent studies have also demonstrated the interaction of UTX with the mitotic deacetylase complex (MiDAC) [12,13]. Interestingly, UTX escapes X-inactivation in females, which is compensated in males by the chromosome Y encoding the *UTY* ortholog (also referred to as *KDM6C*) [14,15]. Although UTY is highly similar to UTX, it displays only weak enzymatic activity [16]. Both proteins play important roles in development and disease with both the enzymatically dependent and independent functions [1,17,18]. For example, comparison of the downstream activities of a catalytically inactive UTX mutant with the wild type (wt) protein in a UTX-deficient BLCA cell line indicated that the tumor suppressor functions of UTX can be enzymatic-independent [19]. The human genome encodes a third human H3K27me3 demethylase JMJD3/KDM6B, which harbors a similar JmjC domain. However, JMJD3/KDM6B lacks the TPR region, and it does not incorporate into MLL3 and MLL4 complexes. In many studies, KDM6 family members plays different and sometimes opposing functions in development [20,21] and in human disease [22]. Many studies have shown the prominent impact of *UTX* mutations on gene expression and diseases. However, whether the mRNA splicing of *UTX* could play a role in its biochemical functions has not been studied so far. Despite identification of different mRNA isoforms in the databases, a comprehensive analysis and quantitative overview of *UTX* alternative splicing events and their distribution over normal and cancerous tissues is lacking.

Here, we focus on a variety of human cell lines and tissue samples to provide of the overall architecture of *UTX* mRNAs. Given the clinical significance of UTX in bladder cancer, we employed long-read sequencing of *UTX* cDNAs from different human cell lines and from normal bladder epithelium or bladder samples to define the alternative splicing region (ASR) of *UTX* mRNAs, which spans exons 12-17 [23]. We expressed the five most abundant *UTX* isoforms to examine their protein functions. The subcellular localization of UTX isoforms is regulated by exon 14 encoding a predicted nuclear localization sequence or NLS. Chromatin association of certain *UTX* isoforms and their protein interactome is controlled by exons located in the ASR.

## Results

### Characterization of the *UTX* alternative splicing events

A comparison of the mRNA isoforms from the human *UTX/KDM6A* gene in various public databases indicated the existence of multiple alternatively spliced mRNAs. The longest isoform (NM_001291415 in the NCBI database) was used in Figure 1A to indicate all possible *UTX* exons. Exon 14, which contains a predicted NLS [24] is absent in the canonical isoform (NM_021140). Therefore, in many human UTX studies an artificial NLS has been attached to canonical UTX to obtain robust nuclear expression, while the long isoform with the natural NLS has been overlooked [7,8,25]. A very similar pattern was observed for *UTY*, with a predicted NLS after the TPR at the long isoform (NM_001258249) encoded by exon 12 (nucleotides 1434-1467), which is absent from the canonical *UTY* transcript (NM_007125) [26].

**Figure 1.**
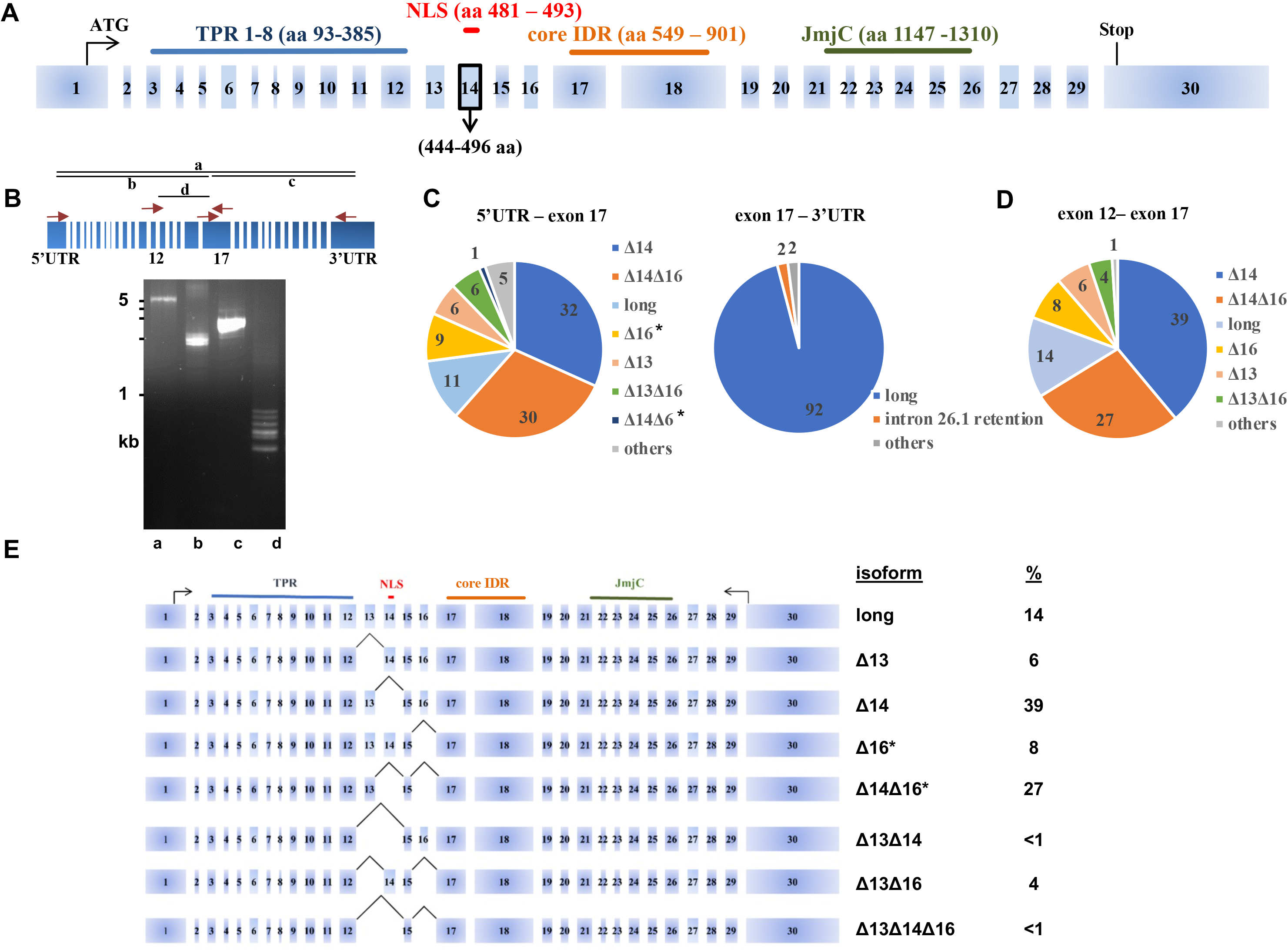
A comprehensive analysis of *UTX* alternative splicing events derived from long-read cDNA sequencing. **A**) Exon-intron structure of the *UTX* gene with the presence of a strong NLS (indicated in red) in the commonly overlooked exon 14. **B**) Different RT-PCR primer sets were used to amplify (parts of) the *UTX* coding region for Nanopore sequencing of RNA isolated from 5637 bladder cancer cells. cDNA products were analyzed by ethidium bromide-staining after electrophoresis on a 1% agarose gel. **C**) Long-read sequencing results of *UTX* cDNAs obtained with primer sets b and c identify the alternative splicing region (ASR) spanning exons 12 through 17. The pie charts indicated the abundance in percentages of the different isoforms in 5637 cells including two novel isoforms Δ16 and Δ14Δ16 (indicated by asterisks). ‘Others’ denote cDNAs of an abundance of 1% or less. **D**) Pie chart of long-read sequencing results of *UTX* cDNAs obtained with primer set d confirmed alternative splicing events between exons 12 and 17 in 5637 cells. Numbers in the pie chart indicate the percentages. **E**) Summary of the results based on the primer-set d are displayed on the *UTX* exon-intron map. Asterisks indicate the novel isoforms.

To investigate the alternative splicing events and the proportion of *UTX* mRNA isoforms in a comprehensive manner, we first focused on the commonly used bladder carcinoma cell line 5637. This epithelial-like cell line has been developed from a grade II urothelial carcinoma of male origin, carrying a wild-type *UTX* allele, and contains no mutations in other members of the COMPASS family of SET1/MLL complexes or other key epigenetic genes [27]. First, we amplified the whole 4359-bp coding region of *UTX* mRNA by RT-PCR (amplicon “a”, Fig. 1B) and subjected the resulting cDNAs to long-read Nanopore sequencing. This experiment revealed a variety of alternative splicing events, which were confined to the central region of the *UTX* gene (data not shown). The hotspot of alternative splicing occurred just after TPR and at the beginning of the middle part of UTX comprising exons 12 to 17. In order to increase the number of high-quality long-reads, we divided the *UTX* coding region into three different regions for RT-PCR amplification of RNAs isolated from 5637 cells. Regions b and c span exons 1 to 17 and exons 17 to 30, respectively, whereas region d covers the central region from exon 12 and exon 17 (Fig. 1B). Agarose gel electrophoresis analysis already revealed distinct DNA fragments of region d cDNAs ranging from 400 to 900 bp (Fig. 1B). We performed long-read sequencing of cDNA products from regions b and, c and we calculated the percentage of different mRNA isoforms of *UTX*. A total of 86,710 reads for region b and 3,898 reads for region c were obtained. Isoforms with less than 100 reads were excluded from further analysis, which was focused on cDNAs with an abundance of 1% or more. The rest were denoted as “others” in Figures 1C and 1D. Analysis of fragment b cDNAs identified many alternative mRNA splicing events in the region spanning exons 12-17 of the *UTX* gene, which we refer to as the alternative splicing region (ASR). Long-read sequencing of fragment c cDNAs, which span the second half of *UTX* (exons 17-30) did not reveal alternative splicing events, except the infrequent retention (2%) of intron 26 encoded sequences (Fig. 1C). These findings are consistent with *UTX* transcripts annotated in the NCBI database, which did not quantify the diversity of *UTX* mRNAs in the ASR. In addition, our analysis identified two novel mRNA isoforms Δ16 and Δ14Δ16, which together comprise more than a third of *UTX* mRNAs (Fig. 1C). Next, we zoomed in by sequencing cDNAs (n=93,817) from region d spanning the ASR (Fig. 1D). Indeed, long-read sequencing confirmed the abundance of two novel *UTX* mRNA isoforms lacking exon 16 as well as isoform Δ14Δ16. To verify the long-read sequencing results we separated individual bands of the ASR-containing fragment d by agarose gel electrophoresis and reamplified the eluted bands for confirmation by standard Sanger sequencing. This confirmed the sequences of the long, Δ14, Δ16 and Δ14Δ16 isoforms (Fig. S1).

In conclusion, *UTX* cDNA analysis from 5637 cells showed that isoform Δ14 is the most abundant mRNA isoform constituting 39% of the *UTX* transcripts followed by the novel isoform Δ14Δ16 with 27%, the “long” isoform (including all 30 exons) with 14%, novel isoform Δ16 with 8%, Δ13 with 6%, Δ13Δ16 with 4%, and all other isoforms at a frequency of less than 1% (Fig. 1D and 1E).

### Long-read analysis of *UTX* mRNAs across a panel of cell lines and bladder samples

Alternative splicing patterns can display both tissue specificity and can be altered in disease states [28]. To examine this issue for *UTX* mRNAs we focused on the *UTX* ASR for mRNA splicing variation across a panel of human cell lines representing different tissues of origin. In total, we performed long-read sequencing on ASR (or fragment d) amplicons from 14 cell lines, three of which with a non-cancer origin and 11 with a cancer origin (including the bladder carcinoma cell lines 5637 and HT-1376). The read numbers per sample range between 11,674 for MCF7 and 125,221 for LNCaP cells (Table S1). The percentage of the longest *UTX* mRNA in 5637 and HT-1376 cells was 14% and 6% (Fig. 2A), respectively, whereas the commonly used *UTX* Δ14 isoform ranged from only 18% in HeLa to 58% in the diploid retinal pigmented epithelial cells (RPE-1). The novel Δ14Δ16 *UTX* mRNA isoform ranged from 19% in RPE-1 to 51% in the colorectal carcinoma cell line, COLO205, whereas the novel Δ16 isoform was less abundant and ranged from 1% to 6% (Fig. 2A).

**Figure 2.**
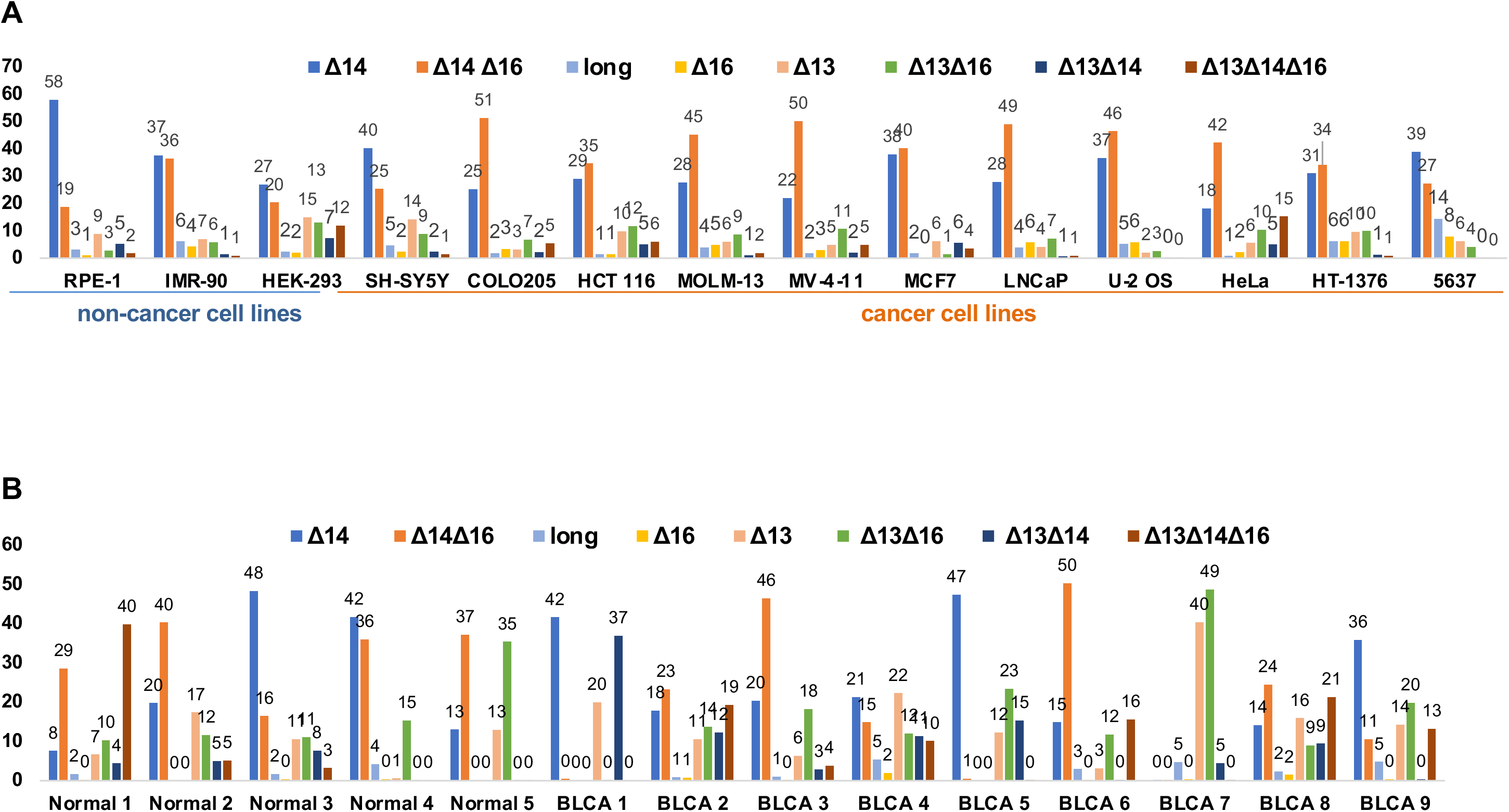
Nanopore sequencing reveals the high frequency of non-canonical isoforms in different cell lines and a high proportion of alternatively spliced transcripts at exon 14 and exon 16. **A)** *UTX* isoform composition in different cell lines was determined by long-read sequencing of *UTX* cDNAs. The frequency of *UTX* isoforms varied between different cell lines. **B)** *UTX* isoform composition in bladder cancer and normal bladder epithelium samples. The frequency of *UTX* isoforms demonstrates a high variability of the isoform distribution in different samples.

Given the high proportion of *UTX* mutations in bladder carcinoma, we decided to investigate the distribution of *UTX* mRNA isoforms in human bladder tissues. For this, we obtained clinical samples from nine BLCAs and from five normal bladder epithelium. All BLCAs were of the muscle-invasive category, and normal epithelial samples were obtained from non-cancer individuals using laser microdissection. Nanopore sequencing of region d spanning the ASR showed a high variability of different isoforms (Fig. 2B). The canonical Δ14 ranged from 8-48% in normal bladder and 14-47% in BLCA, constituting an average of 26% and 24% of the isoforms in normal samples or BLCA, respectively. No significant differences were observed in the proportion of isoforms or exon percentage spliced-in (PSI) values between normal samples and tumors (Table 1). Nevertheless, our results demonstrated a remarkable variability of the *UTX* mRNA isoforms across different samples. The canonical isoform Δ14, which is regarded as the predominant form of *UTX,* makes up only around a quarter of the *UTX* transcripts.

**Table 1.** summary of Nanopopre sequencing results in BLCA and normal bladdeer samples

| Table 1. summary of Nanopopre sequencing results in BLCA and normal bladder samples |  |  |  |  |  |  |  |  |  |  |  |  |  |  |
| --- | --- | --- | --- | --- | --- | --- | --- | --- | --- | --- | --- | --- | --- | --- |
|  | isoform % |  |  |  |  |  |  |  | % of exon skipping |  |  | exon psi |  |  |
| | $\Delta 14$ | $\Delta 14\Delta 16$ | long | $\Delta 16$ | $\Delta 13$ | $\Delta 13\Delta 16$ | $\Delta 13\Delta 14$ | $\Delta 13\Delta 14\Delta 16$ | no ex 13 | no ex 14 | no ex 16 | exon 13 | exon 14 | exon 16 |
| Normal 1 | 8 | 29 | 2 | 0 | 7 | 10 | 4 | 40 | 61 | 81 | 79 | 38 | 19 | 20 |
| Normal 2 | 20 | 40 | 0 | 0 | 17 | 12 | 5 | 5 | 39 | 70 | 57 | 60 | 29 | 42 |
| Normal 3 | 48 | 16 | 2 | 0 | 11 | 11 | 8 | 3 | 33 | 76 | 31 | 67 | 24 | 68 |
| Normal 4 | 42 | 36 | 4 | 0 | 1 | 15 | 0 | 0 | 16 | 78 | 52 | 82 | 21 | 47 |
| Normal 5 | 13 | 37 | 0 | 0 | 13 | 35 | 0 | 0 | 49 | 50 | 73 | 51 | 49 | 26 |
| mean | 26 | 32 | 2 | 0 | 10 | 17 | 3 | 10 | 40 | 71 | 58 | 60 | 28 | 41 |
| BLCA 1 | 42 | 0 | 0 | 0 | 20 | 0 | 37 | 0 | 57 | 79 | 1 | 42 | 20 | 99 |
| BLCA 2 | 18 | 23 | 1 | 1 | 11 | 14 | 12 | 19 | 56 | 73 | 57 | 43 | 26 | 42 |
| BLCA 3 | 20 | 46 | 1 | 0 | 6 | 18 | 3 | 4 | 31 | 74 | 69 | 68 | 26 | 31 |
| BLCA 4 | 21 | 15 | 5 | 2 | 22 | 12 | 11 | 10 | 56 | 58 | 39 | 44 | 42 | 60 |
| BLCA 5 | 47 | 1 | 0 | 0 | 12 | 23 | 15 | 0 | 51 | 63 | 24 | 48 | 36 | 75 |
| BLCA 6 | 15 | 50 | 3 | 0 | 3 | 12 | 0 | 16 | 31 | 81 | 78 | 68 | 18 | 21 |
| BLCA 7 | 0 | 0 | 5 | 0 | 40 | 49 | 5 | 0 | 94 | 5 | 50 | 6 | 94 | 50 |
| BLCA 8 | 14 | 24 | 2 | 2 | 16 | 9 | 9 | 21 | 56 | 69 | 56 | 43 | 29 | 42 |
| BLCA 9 | 36 | 11 | 5 | 0 | 14 | 20 | 0 | 13 | 48 | 60 | 44 | 52 | 39 | 55 |
| mean | 24 | 19 | 3 | 1 | 16 | 17 | 10 | 9 | 53 | 63 | 46 | 46 | 37 | 53 |

In conclusion, we identified two novel mRNA isoforms Δ16 and Δ14Δ16 by applying long-read sequencing of *UTX* cDNAs, which display a high level of variability of splicing isoforms between different cell lines and clinical samples. This variability in mRNA isoforms may instruct different *UTX* functions.

### UTX isoforms display different protein-protein interactions

To compare the protein interactome of the long isoform of UTX with isoforms lacking each of the alternative exons (13, 14 or 16) and the abundant novel UTX isoform Δ14Δ16, we expressed these UTX proteins as N-terminally GFP-tagged versions from a single chromosomal integration site in the Hela-FlpIn/T-REx cell line. The GFP-fusion format allows for both fluorescence localization and interactome experiments for UTX proteins. Immunoblotting of total lysates showed that all GFP-UTX proteins are expressed to similar levels with a slightly lower expression of isoforms Δ13 and Δ16 (Fig. 3A). Confocal fluorescence microscopy showed a robust nuclear expression of GFP-UTX isoforms including exon 14 (Fig. 3B). In contrast, isoforms lacking this exon, Δ14 and Δ14Δ16, also accumulate in the cytoplasm (Fig. 3B), which suggests that the predicted NLS of exon 14 increases nuclear localization of UTX protein. We note in passing that none of the GFP-UTX isoforms expressed in the Hela cells display a punctuated pattern, which would be indicative of biomolecular condensates [7].

**Figure 3.**
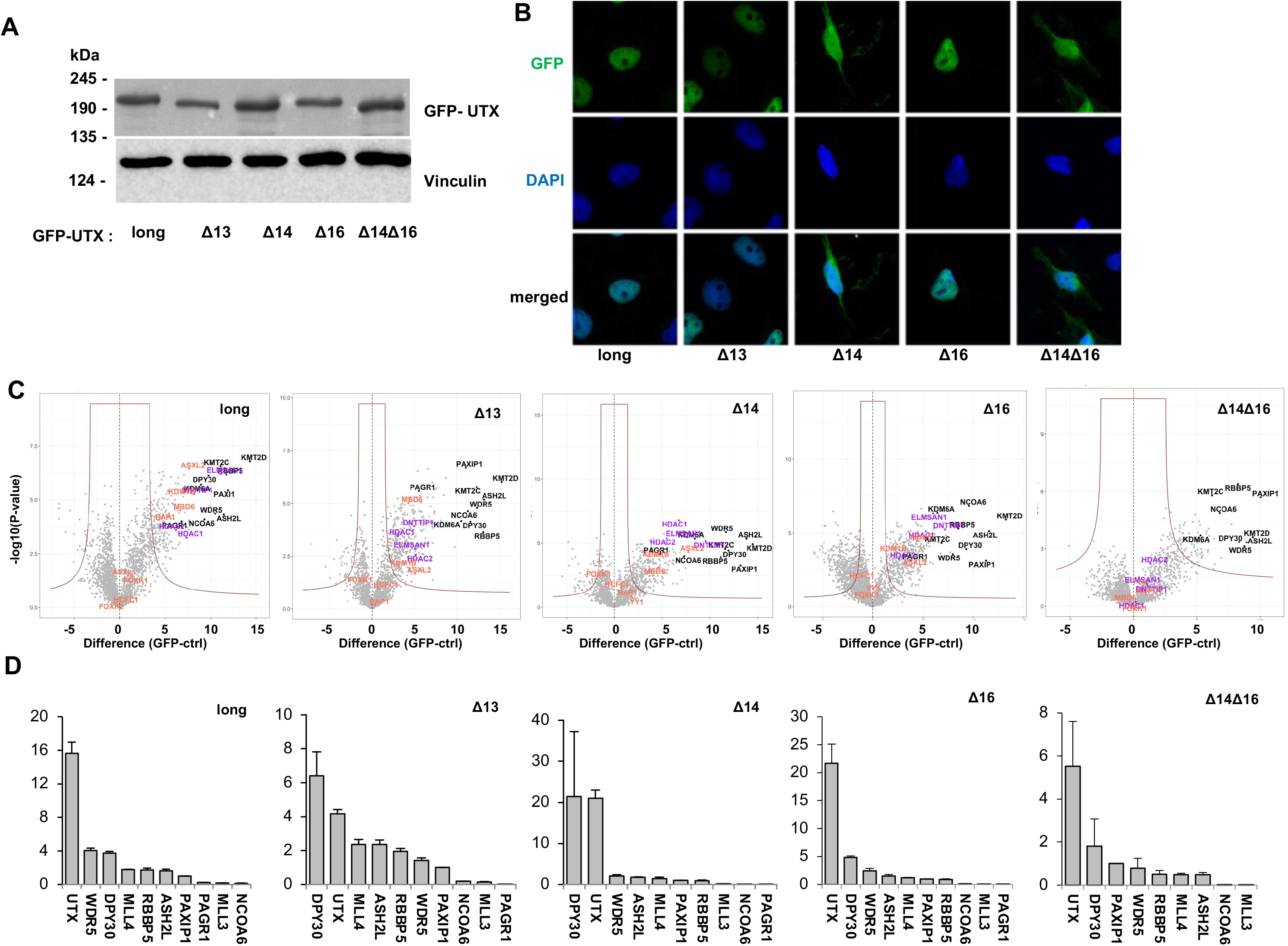
UTX isoforms have different cellular locations and protein-protein interactions. **A**) Immunoblot analysis using anti-GFP antibodies shows that UTX isoforms lacking exon 14 accumulated in the cytoplasm in contrast to isoforms containing exon 14. **B**) Protein-protein interactions of different UTX isoforms as determined by qMS. Interactions with MLL3/MLL4, PR-DUB and MiDAC complexes are marked with black, orange and purple dots, respectively. Isoform Δ14Δ16 does not interact with the MiDAC or PR-DUB complexes. **C**) Comparison of the relative stoichiometries of the MLL3 and MLL4 subunits between UTX isoforms. Protein abundance was normalized to PAXIP1/PTIP peptides. PAGR1 was not detected in isoform Δ14Δ16.

Next, we compared the nuclear interactomes of the five UTX isoforms by conducting GFP-affinity purifications followed by iBAQ-based quantitative mass spectrometry (qMS). Our qMS approach allows both the definition of significant interactors as shown by volcano plots (Fig. 3C) and quantification of the relative abundance of a significant interactor (Fig. 3D) using Perseus software [29]. As expected, [10], we identified the full MLL3 and MLL4 histone H3K4 methylation complexes in the interactome of the long isoform of UTX (indicated by black dots in Fig. 3C). MLL3 and MLL4 subunits also stand out as interactors with all other UTX isoforms indicating that the sequences encoded by exons 13, 14 and 16 are not essential for MLL3/MLL4 interactions, which is confirmed by their relative stoichiometries normalized against the PAXIP1 subunit (Fig. 3D). UTX seems to interact stronger with the MLL4 compared to the MLL3 complex, but this may be related to the relative expression levels of MLL3 and MLL4 in HeLa cells [10]. The NCOA6 and PAGR1 proteins are sub-stoichiometric subunits of MLL3/MLL4 complexes as we reported before [10]. In addition, and with lower stoichiometries (Fig. 3C, S2A and S2B), we observed members of the PR-DUB H2A deubiquitinase [30] and MiDAC histone deacetylase [31] complexes as significant hits in the UTX interactomes. All four MiDAC subunits DNTTIP, ELMSAN1, HDAC1 and HDAC2 are present in GFP-purifications of UTX long, Δ13, Δ14 and Δ16 (Fig. 3C and S2A). Interestingly, MiDAC subunits were completely absent from the UTX Δ14Δ16 interactome. Several but not all PR-DUB members (BAP1, MBD6, KDM1B, and ASXL2) were present in the interactomes of UTX long, Δ13, Δ14 and Δ16 (Fig. 3C, S2A and S2B). Whereas the BAP1 catalytic subunit of PR-DUB was identified as a significant interactor of UTX long, it was absent with the other isoforms (Fig. 3C and S2B). This indicates that PR-DUB interactions are more sensitive to the absence of UTX exons 13, 14 or 16, which is in contrast to MiDAC interactions. Interestingly, both MiDAC and PR-DUB subunits were not identified with the Δ14Δ16 isoform of UTX (Fig. 3C). This indicates that the combination of sequences encoded by exon 14 and 16 of UTX are involved in interactions with the PR-DUB and MiDAC histone modification complexes.

Taken together, we determined the nuclear interactors for five abundant UTX isoforms to find that they all efficiently incorporate into the MLL3 and MLL4 complexes. In addition, UTX proteins interacts with members of the PR-DUB and MiDAC histone modification complexes at lower stoichiometries, and these interactions are sensitive to loss of ASR exons.

### The combination of middle part with TPR is required for proper protein-protein interaction of UTX

In order to better define the UTX regions important for the observed protein-protein interactions we expressed GFP-tagged UTX fragments in Hela cells to perform qMS (Fig. 4A). First, we focused on the middle part of UTX (excluding TPR repeats and JmjC domain and covering residues 398-932, which spans exons 12 to 18 and includes the ASR. Interestingly, the GFP fusion of this middle part did not interact with any subunit of the MLL3, MLL4, MiDAC or PR-DUB complexes (Fig. 4B). When combining the TPR with the middle part (residues 2-880), we observed all subunits of the above-mentioned complexes as significant interactors (Fig. 4C). In addition, the middle part plus JmjC (residues 398-1453) interacts with the MLL3 and MLL4 complexes at a low stoichiometries, when compared to the TPR with the middle part protein (Fig. 4C, 4D and S4). Importantly, the middle part plus JmjC of UTX does not display interactions with subunits of the PR-DUB or MiDAC complexes. Taken together, these results indicate that the middle region of UTX spanning the ASR is not able to interact independently with the subunits of the MLL3/MLL4, MiDAC or PR-DUB complexes. This region possibly stabilizes these interactions in combination with the TPR of UTX. Both the TPR and JmjC domains can mediate interactions with the MLL3 and MLL4 complexes, but the TPR seems the predominant interaction region for MiDAC and PR-DUB.

**Figure 4.**
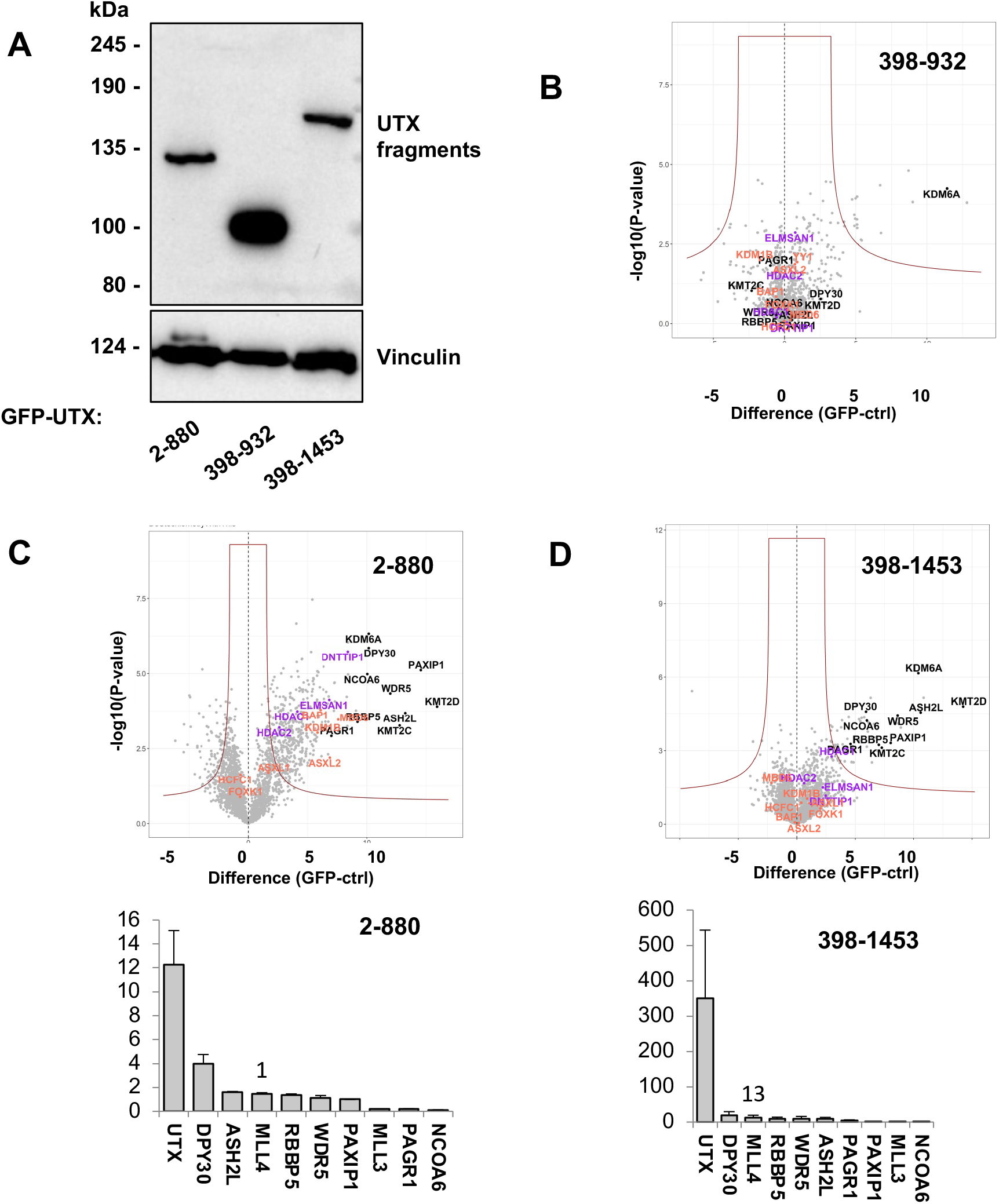
Protein interaction analysis at GFP-tagged UTX fragments. **A**) Immunoblot analysis of GFP-tagged UTX fragments expressed in HeLa-FRT cells for qMS analysis. **B**) UTX 398-880 (middle part) including ASR did not pull down any members of MLL3/MLL4, PR-DUB or MiDAC complexes. **C**) UTX 2-880 (TPR and middle part) including ASR expression identified all subunits of the MLL3/MLL4, PR-DUB and MiDAC complexes. UTX 398-1453 (middle part combined with JmjC) showed low stoichiometry interactions of the MLL3 and MLL4 complex subunits and does not interact with subunits of the PR-DUB or MiDAC complexes.

### The UTX long isoform displays a stronger chromatin association when compared to canonical isoform **Δ**14

In order to examine the effect of exon 14 sequences on the genome localization properties of UTX proteins, we applied greenCUT&RUN profiling [32] for UTX long and Δ14 in Hela-FRT cells, that exon 14 is missing from most of its spliced transcripts (Fig. 2A). The distribution of binding over different functional genomic regions was similar between the two isoforms (Fig. 5A). However, and as expected from the increased nuclear abundance, a higher number of peaks were observed for UTX long compared to Δ14 (n=14,950 and n=9,070, respectively; Fig. 5B). The heatmaps of UTX long and UTX Δ14 (Fig. 5B) show that peaks identified with the UTX long isoform were also identified with the Δ14 isoform. Comparison of the chromatin properties of UTX bound regions for the histone marks H3K4me1, H3K27ac, H3K27me3, and ATAC sites did not reveal clear differences between the UTX long and Δ14 isoforms (Fig. 5C) For both isoforms we observed robust binding to active chromatin sites as indicated by the overlap with H3K4me1, H3K27ac and ATAC signals (Fig. 5E). We note that H3K27me3 sites are under-represented at UTX sites (Fig. 5C). Next, we examined whether the two isoforms display differential genomic binding by selecting regions of significant changes. In total, only eleven differential sites were identified using default parameters (FC of > 4 and p-value < 0.0001) (Fig. 5D and Table S2). Genomic tracks for the three representative UTX long-specific sites are shown in Figure S4, which indicates that no or fewer reads were present with UTX Δ14. This suggests that these regions are sensitive to levels of nuclear UTX. However, these regions do not map to known gene promoters or enhancers casting uncertainty of their relevance for gene control. In conclusion, the genome localization analysis indicates that exon 14 sequences do not control specific chromatin binding properties of UTX, but rather its nuclear abundance, which is reflected by the lower peak number of UTX Δ14. Both UTX long and Δ14 isoforms bind to the active regions of the genome as revealed by colocalization with histone H3K4me1 and H3K27ac modifications and with ATAC signals.

**Figure 5.**
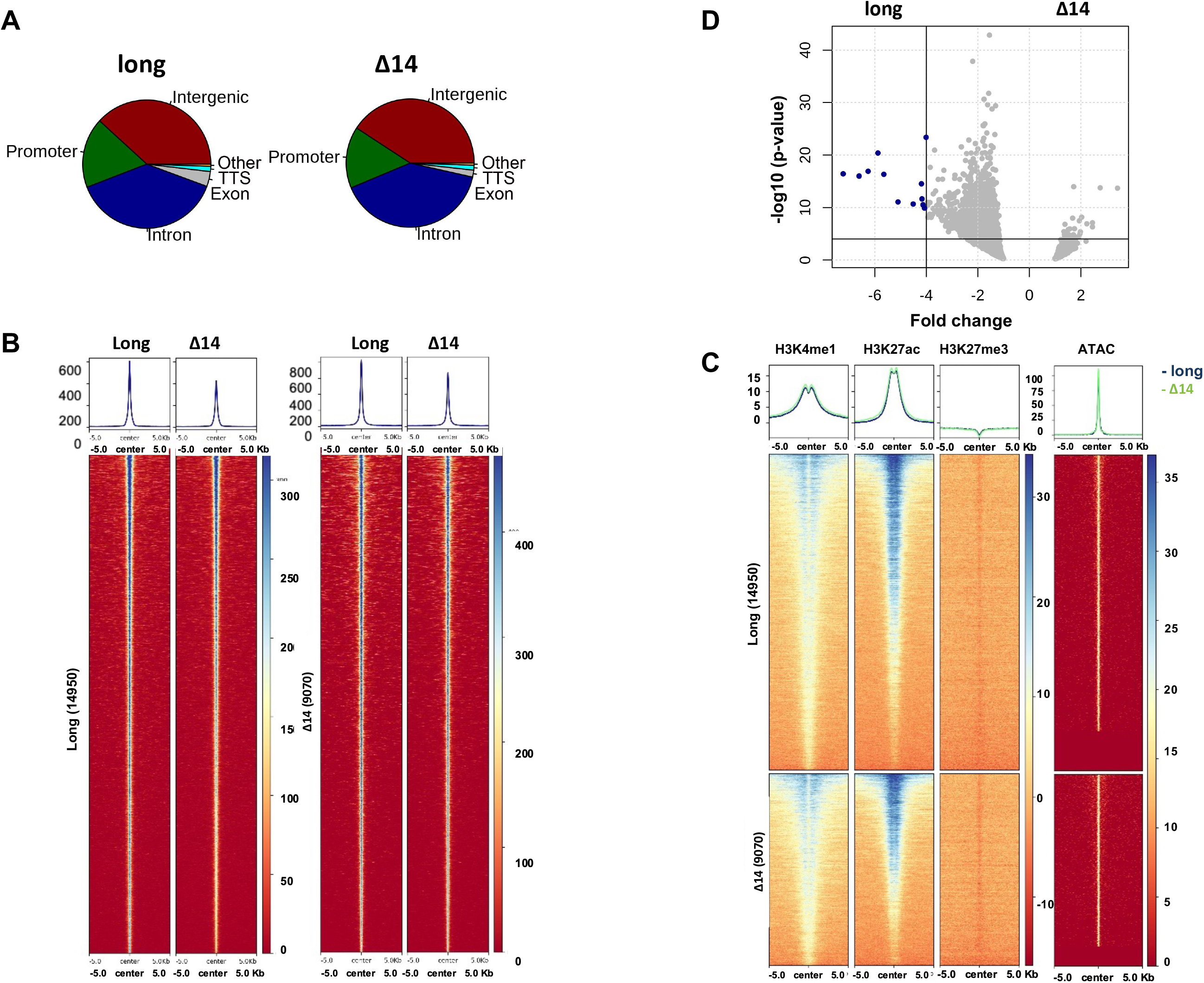
GreenCUT&RUN analysis for UTX long and Δ14 shows stronger chromatin binding by the long isoform, while both isoforms are equally enriched at the relevant histone modification sites. **A)** The ratio of UTX binding to different genomic regions for isoforms long or Δ14. **B)** Heatmaps show the higher coverage for UTX long isoform. The total number of peaks was 14,950 for UTX long and 9,070 for Δ14. **C)** Heatmaps show no difference for H3K4me1, H3K27ac, and H3K27me3 at the binding sites of either UTX isoform. UTX peaks are overlapping the ATAC-seq signals, which showed that UTX binds to the active regions of chromatin, regardless of the isoform. **D)** V-plot demonstrating significant genes for the long isoform compared to Δ14.

## Discussion

Next-generation sequencing data of healthy and pathological samples, especially from cancer tissues have indicated that *UTX/KDM6A* is an important gene for human diseases [1]. Molecular studies revealed that the UTX H3K27 demethylase plays a pivotal role in gene expression control as a regulator of chromatin modifications. The molecular functions of UTX are believed to be related to its interaction with the MLL3 and MLL4 complexes [11,33], which involves the TPR domain of UTX. Besides the importance of TPR and the catalytic JmjC domains, the middle part of UTX contains an intrinsically disordered domain (IDR), which has been implicated recently in the formation of biomolecular condensates [7]. In this study we examined the alternative splicing patterns of UTX mRNAs across a variety of human cell lines and bladder tissue samples. It is well known that alternative splicing provides an additional layer of complexity to the diversity of protein functions and interactions. More than 90% of human genes are subjected to alternative splicing isoforms [34]. Different protein isoforms may play different and sometimes opposing roles [35]. Analysis of HAVANA annotated transcripts has shown that alternative splicing affects the domain architecture in around 43% of the genes [36]. The tetratricopeptide repeat superfamily was the fourth top domain to be affected by domain architecture difference between splicing isoforms.

In this study we discovered the presence of an alternative splicing region (ASR) in *UTX* mRNAs in the middle part of the protein, which does not overlap with the core IDR (Fig. 1A). Exons 13, 14 and 16 frequently alternated in *UTX* mRNAs. As a result, a variety of different UTX mRNA splicing patterns were observed in normal bladder, BLCA and different cell lines. The frequency of these isoforms was variable and the canonical isoform lacking exon 14 comprised only a quarter of all isoforms on average. The alternative splicing of exons 14 and 16 is especially interesting, since this region is involved in protein-protein interactions of UTX with PR-DUB and MiDAC complexes (Fig. 4).

For exon 14, considering its relative abundance, the presence of a predicted NLS was associated with an increased nuclear localization of the corresponding UTX isoform. Indeed, the absence of exon 14 correlates with a reduced genomic binding of UTX, while the proteins are expressed to similar levels in the cells. Since its cloning, the weak nuclear expression of an important histone modifier such as UTX has puzzled researchers [37]. To solve the issue, the ORF of UTX has been fused with an artificial NLS in many studies, ever since [8]. We have solved this paradox here, by describing different cellular distributions for distinct UTX isoforms. Isoforms including the NLS-encoding exon 14 should be under further focus, because it is possible that the main chromatin functions of endogenous UTX are attributed to these isoforms rather than cytoplasmic UTX. The *UTX* Δ14 isoform has been considered as the reference in different clinical studies. For this reason, there is no mutation analysis for UTX exon 14 in the TCGA dataset and cBioportal [38]. We found in the International Cancer Genome Consortium database that exon 14 harbors cancer mutations, such as the recurrent mutations at the chrX:45060619 locus (GRCh38, Fig. S5). We also examined the cytoplasmic fraction of the GFP-UTX isoforms in qMS experiments and we observe only weak interactions with some of the MLL3/MLL4 complex members (data not shown). On the other hand, it is possible that there are specific cytoplasmic functions or there is a dynamic equilibrium between the cytoplasmic and nuclear localization of the abundant isoforms lacking exon 14, which may depend on its interaction with the MLL3 and MLL4 complexes [6].

In our study we identified novel isoforms of *UTX* mRNAs lacking exon 16. The observation of UTX interactions with PR-DUB and MiDAC complexes is interesting as it provides a mechanistic basis for histone modification crosstalk orchestrated by UTX. We speculate that the PR-DUB and MiDAC complexes add a regulatory layer to gene expression regulation by UTX proteins. Moreover, a different interaction spectrum of UTX isoform Δ14Δ16 was of particular importance. A special regulatory role for different UTX isoforms could be presumed that are regulated at a splicing level. The MiDAC complex has been suggested to recruit HDAC1 and HDAC2 to deacetylate H3K27ac and H4K20ac [13]. We found differences in chromatin binding between UTX long and Δ14, which may relate to difference in nuclear abundance of these isoforms. As expected UTX long is the nuclear specific isoform and probably most of chromatin-related functions of UTX are attributed to this and other exon 14-including isoforms. Especially, the considerable abundance of these isoforms as revealed by our Nanopore results indicates the importance of alternative splicing for differential UTX functions in general and for overall UTX mRNA abundance levels in different tissues and cell types, in particular. Development of isoform-specific antibodies and cancer mutation analysis in the alternative exons are among such new directions for further analysis.

Splicing factor mutation is one of the processes implicated in tumorigenesis [39]. Bladder carcinoma and uveal melanoma show the highest ratio of splice factor mutations among TCGA cancers [40]. Interestingly, UTX is amongst the most frequently mutated tumor suppressors in these cancer types [2]. It is tempting to speculate that the observed diversity in *UTX* mRNAs is related to the overall differences in splicing factor activity in the cell types and tissue samples analyzed in our study. These observations motivate future studies linking splicing factor mutations to *UTX* mRNA splicing events in cancer-susceptible tissues like the bladder.

In conclusion, this study provides a detailed analysis of the architecture of *UTX* mRNAs in several cell types. Alternative splicing events of *UTX* pre-mRNAs are localized to exons 12 to 17, which are involved in the nuclear localization of UTX proteins and their interactions with the PR-DUB and MiDAC chromatin regulatory complexes.

## Materials & Methods

### Clinical samples, cell lines and cloning

Clinical material was provided with allowance of the RWTH centralized Biomaterial (RWTH cBMB), Medical Faculty, RWTH Aachen University. Patients’ consents were obtained before conducting the study. This study was conducted in accordance with the Declaration of Helsinki. The local ethical committee approved the protocol for this study.

The *UTX* long isoform cDNA was purchased from Genscript (CloneID OHu24601). The cDNA for *UTX* lacking exon 14 sequences was PCR amplified from pCMV-HA-UTX, which was a gift from Kristian Helin (Addgene plasmid # 24168) [8]. All UTX isoforms and mutants were generated by PCR cloning into the pDONR221 vector or by site-directed mutagenesis using the QuikChange strategy (Agilent, CA, USA). Inserts were transferred to the pCDNA5_FRT_TO_N-GFP destination vector using GATEWAY cloning according to instructions of the manufacturer (Life technologies, Thermo Fisher Scientific, MA, USA). All plasmids were validated by DNA sequence analysis and primer sequences are available upon request.

Hela FlpIn/TRex cells were stably transfected with pOG44 and pCDNA5-based vectors for expression of GFP-tagged UTX and were selected with blasticidin and hygromycin as described before [10,41]. RPE-1 cells were grown in DMEM: F-12 Medium (1:1), IMR-90, SH-SY5Y, MCF7, and HT-1376 in Eagle’s Minimum Essential Medium (EMEM), HEK293 and Hela in DMEM, COLO205, MV-4-11, Molm13, LNCaP and 5637 in RPMI-1640 Medium, HCT 116, and U-2 OS in McCoy’s 5A. All media were supplemented with 10% FBS.

### Immunofluorescence and confocal microscopy

Hela cells carrying GFP-tagged UTX isoforms were induced 24 hours with 1 µg/ml doxycycline before the experiment. The cells were fixed in 4% formaldehyde in PBS, permeabilized with 0.1% Triton X-100 in PBS, and blocked with 10% normal goat serum in PBS. DNA was stained with DAPI, and slides were imaged with ZEISS LSM880 Airyscan. ZEISS ZEN 2.3 software was used for image analysis.

### Immunoblotting analyses

To prepare whole cell lysates, cells were lysed in Laemmli Buffer with 50 µg/ml DTT. Lysates were incubated at 95°C for 10 min to denature the proteins before separation by SDS-polyacrylamide gel electrophoresis. Proteins were transferred to the nitrocellulose membrane (Invitrogen) by electroblotting. The membrane was blocked in 5% skimmed milk and incubated overnight at 4°C with primary antibodies for UTX (Cell Signaling Technologies #33510), GFP (JL-8, Clontech) or Vinculin (7F9, SantaCruz) as a loading control. Blots were incubated with corresponding secondary antibodies at concentrations suggested by the manufacturer (Bio-Rad), were developed using the Clarity Western ECL kit (Bio-Rad, Hercules, CA) and were imaged using a ChemiDoc Touch system (Bio-Rad Hercules, CA).

### GFP affinity purification and MS sample preparation

HeLa FlpIn/TREx cells carrying the doxycycline-inducible GFP-UTX allele were treated for 48 h with 1 µg/ml doxycycline. GFP expression was verified using immunoblotting using GFP (JL-8, Clontech) and α-tubulin (CP06, Calbiochem) antibodies. Nuclear and cytoplasmic extracts were prepared for GFP-affinity purification coupled to mass spectrometry analyses as described before (8). In brief, about 300 million cells were harvested after induction with 1 µg/mL doxycycline for 48 h, washed twice with PBS (Gibco, #10010-015), was resuspended in 5 volumes of cold Buffer A (10 mM Hepes-KOH pH 7.9, 1.5 mM MgCl_2_, 10 mM KCl), and incubated for 10 min on ice. The cells were pelleted and resuspended in 2 volumes of Buffer A supplemented with 1 µM DTT, 0.5 % NP-40 and cOmplete proteinase inhibitor (CPI, Roche, #11836145001, referred to as buffer A complete, hereafter). To separate nuclear and cytoplasmic lysates, the cells were then homogenized in a Dounce homogenizer, on ice. The nuclear fraction was pelleted by centrifugation at 3,300 *g* for 15 minutes at 4°C. The supernatant was further cleared from debris by centrifugation at 16,000 g and 4°C for 1 h, and further processed as cytoplasmic fraction. The nuclear pellet was then washed out of the cytoplasmic carryover by adding 10× volume buffer A complete and centrifugation at 3,300 g for 5 minutes. The pellet was then resuspended and gently agitated in high salt Buffer B (420 mM NaCl, 20 mM Hepes-KOH pH 7.9, 20% v/v glycerol, 2 mM MgCl_2_, 0.2 mM EDTA, 0.1 % NP40, 1x CPI, 0.5 mM DTT) at 4°C for 1.5 h. Subsequently, the supernatant representing the nuclear extract was obtained by centrifugation at 16,000 *g* and 4°C for 1 h.

After Bradford protein measurement, 1 mg of nuclear and 2 mg of the cytoplasmic fraction were used for GFP, or control pulldowns as described before [42]. GFP-coated agarose beads (Chromotek) or control agarose beads (Chromotek) were added to the protein lysates in three replicates each and rotated overnight at 4°C in binding buffer (20 mM Hepes-KOH pH 7.9, 300 mM NaCl, 20% glycerol, 2 mM MgCl_2_, 0.2 mM EDTA, 0.1% NP-40, 0.5 mM DTT and 1x CPI). Thereafter, the beads were washed twice with the binding buffer containing 0.5% NP-40, twice with PBS containing 0.5% NP-40, and twice with PBS. On-bead digestion of bound proteins was performed overnight in elution buffer (100 mM Tris-HCl pH 7.5, 2 M urea, 10 mM DTT) with 0.1 µg/ml of trypsin at RT and eluted tryptic peptides were bound to C18 stage tips (ThermoFischer, USA) prior to mass spectrometry analysis.

### Quantitative mass spectrometry analysis

Samples were analyzed by nanoflow-LC-MS/MS on a Q-Exactive Plus coupled to an Easy-nLC 1000 or an Orbitrap Fusion Lumos coupled to an Easy-nLC 1200 nanoflow-LC-MS system (ThermoFisher Scientific). A flow rate of 300 nl/min and a gradient of increasing organic proportion (buffer A: 0.1 % formic acid, buffer B: 0.1 % formic acid in 80 % acetonitrile) in combination with a reversed phase C18 separating column of 25 cm length was used for peptide separation. Each MS scan was followed by a maximum of 10 MS/MS scans in the data-dependent mode (TOP-10 method). Blanks were run between sample sets (e.g. between GFP and agarose control sample sets).The outcome raw files were analyzed with MaxQuant software (version 1.5.3.30). Data were aligned to Uniprot human FASTA database [43]. Volcano plots were generated using Perseus (MQ package, version 1.5.4.0). Contaminants, reverse peptides, and protein identification based on only one replication were filtered from raw data. Label-free quantification (LFQ) values were transformed to the log2 scale to generate the normal distribution of the data. Quality was checked by generating the unsupervised clustering of replicates and predicted proteins that were depicted as a heatmap for manual inspection. Scatter plots of the hits were also generated based on the Spearman’s correlation coefficient of the LFQ values to quality check the correlation between the GFP condition of each experiment. Imputation of the missing values was then performed on the normally distributed data (width= 0.3 and shift= 1.8). The significantly different proteins between GFP and agarose control pulldown proteins were calculated using a two-tailed Student’s t-test using 1% FDR. The constant value of 1 was kept for the threshold of significance (S0=1). Intensity Based Absolute Quantification (iBAQ) values were used to calculate the stoichiometry as the subsequent relative protein abundance estimation [44]. The iBAQ values for each replication of the GFP pulldown were subtracted by the mean of the values from agarose bead control pulldowns. The abundance of nuclear interactors was normalized based on the PAXIP1 subunit of the MLL3 and MLL4 complexes.

### RT-PCR, gel electrophoresis, and Nanopore analysis of the UTX isoforms

For cell lines RNA was extracted using the RNeasy kit (Qiagen), and for the clinical samples the truXTRAC FFPE kit was used (Covaris). DNase treatment was performed using the Turbo DNase kit (ThermoFisher). cDNA was generated with SuperScript III (ThermoFisher) and polyT primers. TAKARA DNA polymerase (TAKARA) was used for PCR reactions. The entire UTX ORF was amplified using primers targeting 5’-UTR and 3’-UTR (Table S3) and analyzed by ethidium bromide staining after separation using a 1% agarose gel. The ASR region was amplified using primers targeting exon 12 and exon 17 of *UTX*. Observed bands were gel eluted, reamplified, cleaned up and subjected to Sanger sequencing.

The primers also included an extension of Nanopore universal tags:

5’-TTTCTGTTGGTGCTGATATTGC-[project-specific forward primer sequence]-3’

5’-ACTTGCCTGTCGCTCTATCTTC-[project-specific reverse primer sequence]-3’.

For Nanopore sequencing, PCR products were further processed following the manufacturer’s instructions (PCR barcoding 96 amplicons SQK-LSK109). 0.5 nM of the first-round PCR was further amplified using the Nanopore Barcoding primers (EXP-PBC096). Thereafter, the barcoded amplicons were pooled, and a 0.75x AMPure bead clean-up (A63880, Beckman Coulter) performed to deplete unwanted fragments below 150 bp. Thereafter, the pool was subjected to NEBNext FFPE DNA Repair and Ultra II End-prep kits (M6630 and E7546, New England Biolabs). After that, Nanopore adaptors were ligated using NEBNext Quick T4 DNA ligase (E6056, New England Biolabs) and subsequently cleaned with AMPure XP beads (A63880, Beckman Coulter) using Nanopore’s short fragment buffer for washing the beads. The library was loaded on MinION Flow cells (FLO-MIN106, Oxford Nanopore) using the supplied Sequencing Buffer and Loading Beads. Raw data was basecalled using Guppy (version 4.3.2). Analysis was performed using FLAIR software (version 1.5.1) (https://github.com/BrooksLabUCSC/flair). In brief, reads were mapped to human genome (version hg38) using minimap2 with option: -ax splice -t 30 --secondary=no. Each aligned bam files were converted to bed file using bam2Bed12.py tool of FLAIR and then misaligned splice-sites were corrected with genome annotation available at GENCODE project’s website (https://www.gencodegenes.org/) (version 32). In order to identify all highly confident isoforms present in cohort, splice-site corrected data from all samples were pooled together. After this, collapse function of FLAIR was used to merge identical isoforms. Reads associated with these collapsed isoforms were quantified for each sample separately. Isoforms with read number of less than 100 or with an occurrence of < 1% were filtered out.

### Public dataset analysis

NLS prediction was performed using the NLS mapper tool [24]. The mutation information was collected from the cBioPortal and gnomAD web browsers [40,45,46]

### GreenCUT&RUN

Half a million Hela-FRT cells expressing either UTX long or Δ14 isoforms as GFP fusions were harvested and washed twice in cold PBS. The cells were immobilized on concanavalin A-conjugated paramagnetic beads, permeabilized with 0.05% digitonin and subjected to greenCUT&RUN protocol as described before [47]. We added mononucleosomal Drosophila DNA as spike-in DNA for normalization purposes. Sequencing libraries were prepared using NEB Next Ultra II kit (New England Biolabs). The resulting DNA quantity and size distribution was assessed by Qubit instrument (Invitrogen, USA) and Agilent Bioanalyzer chips (DNA high sensitivity assay), respectively.

### Bioinformatic analyses of greenCUT&RUN

The HeLa cell data for H3K4me3 (ID: ENCFF063XTI), H3K4me1 (ENCFF617YCQ), H3K27ac (ENCFF113QJM) were obtained from ENCODE consortium (https://www.encodeproject.org/) and ATAC-seq from SRA-NCBI (accession ID: SRR8171284). Quality control filtering was performed using Trim-galore (version 0.6.3) with default parameters. The good quality reads were aligned on the human (version hg38) and Drosophila reference genome (BDFP5) using bowtie2 (version 2.3.4.1) with option: – dovetail –local –very-sensitive-local –no-unal –no-mixed –no-discordant -I 10 -X 700 [32,48]. We used HOMER for calling the narrow peaks with default parameters except, disabled “filtering based on clonal signals” with option: -C 0. The peaks were annotated with HOMER. Those peaks present in the miRNA, ncRNA, pseudogenes, snoRNA, scRNA and rRNA were categorized as “other”. To generate Heatmaps, computeMatrix function of the deeptools (version 3.3.2) was used with default parameters except sum of the reads were calculated per bin using option “--averageTypeBins” instead of mean. To normalize this matrix, sum of the reads were divided either by total number of the reads (if not available e.g. ChIPseq data downloaded from ENCODE website) or by SpikeIn reads. To generate heatmaps for the ENCODE datasets bamCompare files were used.

## Data availability

The greenCUT&RUN and Nanopore datasets have been deposited to the Sequence Read Archive (SRA) portal of the NCBI with Bioproject accession ID PRJNA952530. All mass spectrometry data were deposited to Xchange Proteome with accession number PDX041254.

## Supporting information

This article contains supporting information.

## Acknowledgements

The authors would like to acknowledge the work of Dr. Roy Baas for his preliminary mass spectrometry investigations on UTX. We also thank Prof. Michiel Vermeulen (Radboud Institute for Molecular Life Sciences, Radboud University Nijmegen) help and advice with mass spectrometry in the initial phase of the project. We thank Dr. Christoph Schell (Institute of Surgical Pathology, Medical Centre – University of Freiburg) for providing help with the initial clinical samples. We are grateful to Prof. Christoph Peters (Institute for Molecular Medicine and Cell Research, University of Freiburg) for support. We appreciate the assistance of Ms. Ursula Schneider with preparing the clinical samples. We gratefully acknowledge the support of the Bioinformatics facility of the Max-Planck Institute of Immunobiology and Epigenetics. We thank Timothy Chan and Ezgi Özerli-Gøknar for critical reading of the manuscript. We also thank all members of the Timmers lab and especially Timothy Chan, Md Saiful Islam and Simona Capponi for discussions and support.

## Author contributions

OF conducted the experiments. OF and SN contributed to acquisition of data and analysis. OF, SN and MT contributed to the design, conception, and interpretation of data. SF contributed in Nanopore. MB contributed to MS experiments. RKC contributed to provide clinical samples and advice. All authors contributed to conception, drafting the article and revisions in intellectual content and approved it for publication.

## Funding and additional information

This research was financially supported by the grants from the Deutsche Forschungsgemeinschaft (DFG, German Research Foundation) by 192904750-SFB 992, and SFB850 subproject B9.

## Legends of Supporting Information

**Figure S1. Confirmation of the long-read sequencing results by standard Sanger sequencing.**

Agarose gel analysis and extraction of four individual *UTX* cDNA fragments from 5637 cells for re-amplification. **B**) Sanger sequencing results of gel extracted bands. Multiple sequence alignment indicating the most abundant isoforms, which confirms the Nanopore sequencing results.

**Figure S2. Stoichiometries of non-MLL3/4 subunit interactors of UTX isoforms.**

**A**) Relative stoichiometries of the MiDAC subunits for the different UTX isoforms are normalized against the MLL3/MLL4 subunit PAXIP1. UTX Δ14Δ16 did not show any significant interactions with MiDAC subunits. **B**) Relative stoichiometries of the PR-DUB subunits for the different UTX isoforms are normalized against the MLL3/MLL4 subunit PAXIP1. The UTX Δ14Δ16 isoform did not show any significant interactions with PR-DUB subunits. BAP1 was not detected with the Δ13, Δ14, Δ16 isoforms, and ASXL1 was not detected with Δ16 isoform.

**Figure S3. PR-DUB and DNTTIP1 complex stoichiometry of the qMS for UTX fragments.**

**A**) The stoichiometry for MiDAC complex subunits that were identified with UTX fragment 2-880 (TPR + middle part). HDAC2 was not detected. **B**) The stoichiometry for PR-DUB complex that were identified with UTX fragment 2-880 (TPR + middle part).

**Figure S4. Representative IGV tracks**

UTX peaks that are more strongly bound by the long isoform compared to the Δ14 isoform at figure 5 are shown at the *TTC1* (**A**) *GLUD2* (**B**) loci.

**Figure S5. *UTX* exon 14 mutations in the ICGC and GnomAD databases for genetic mutations.**

The overlooked *UTX* mutations located in exon 14. Bold locations have been repeated in different samples. Asterisk indicates a mutation in the predicted NLS.

