## Supplementary figures and images for "Alternative mRNA splicing controls the functions of the histone H3K27 demethylase UTX/KDM6A"

### Figure S1

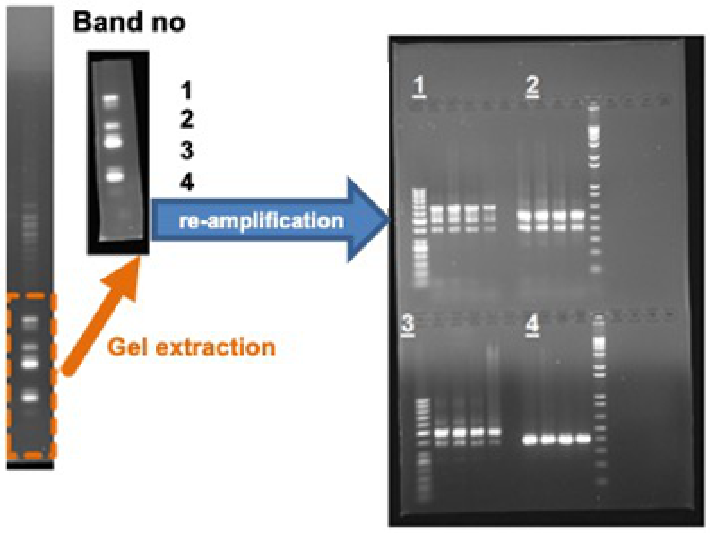

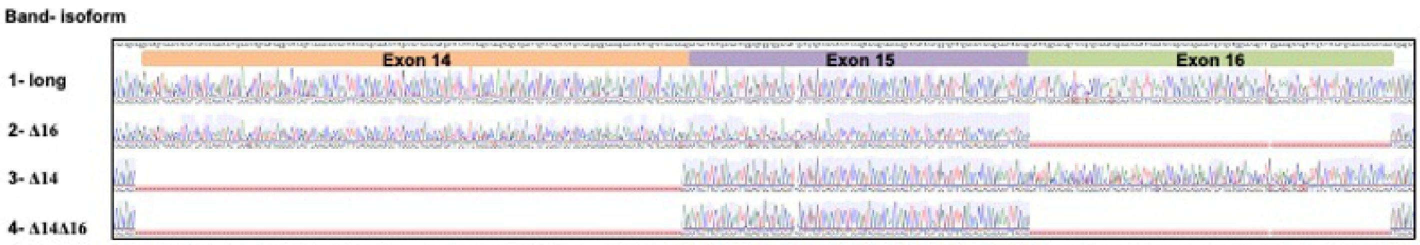

### Figure S2

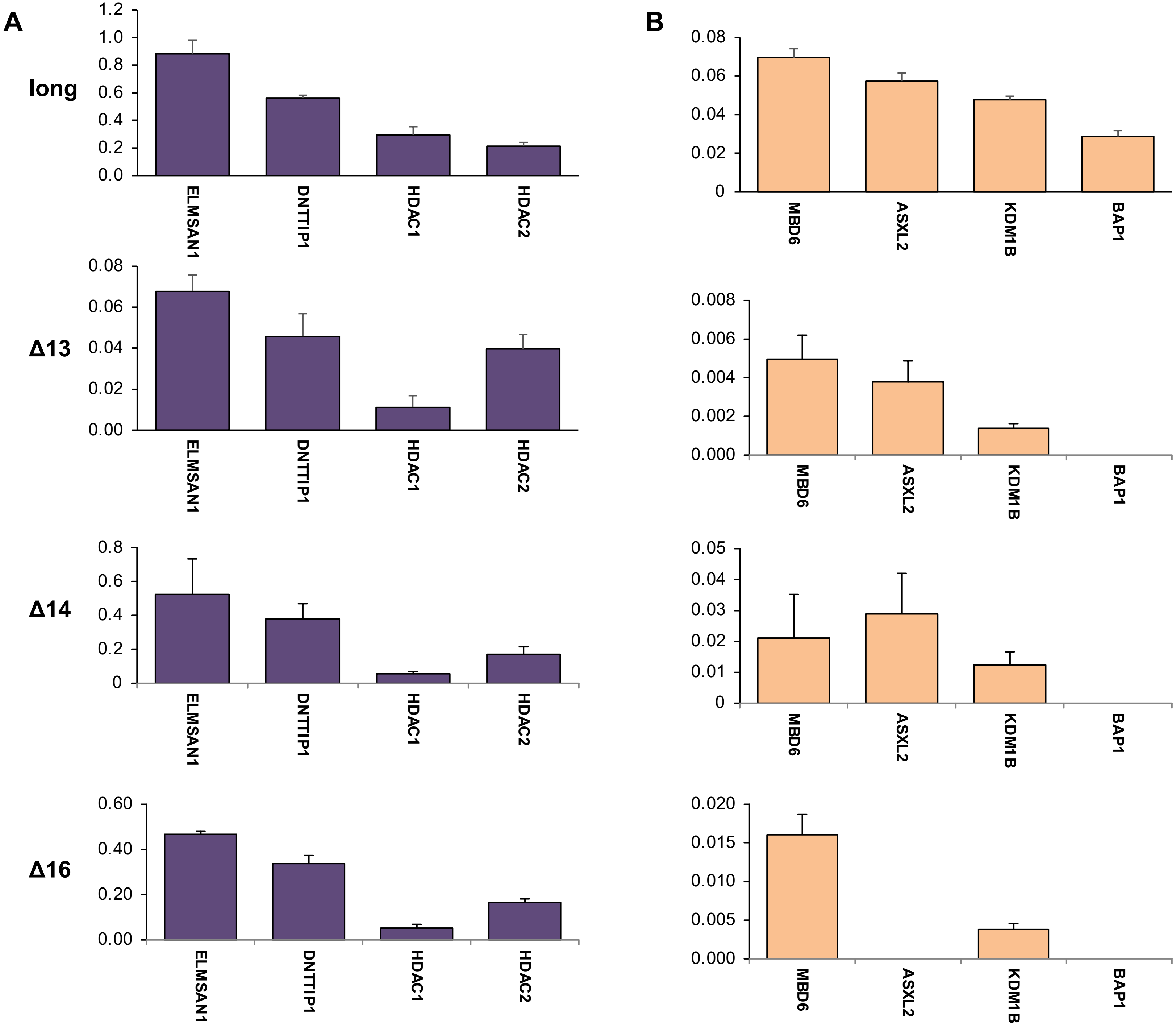

### Figure S3

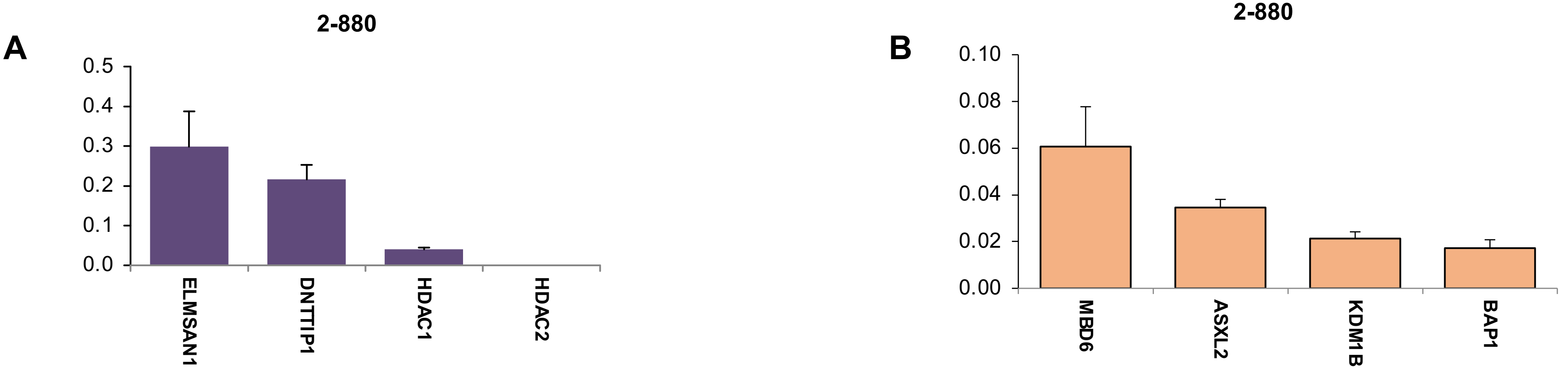

### Figure S4

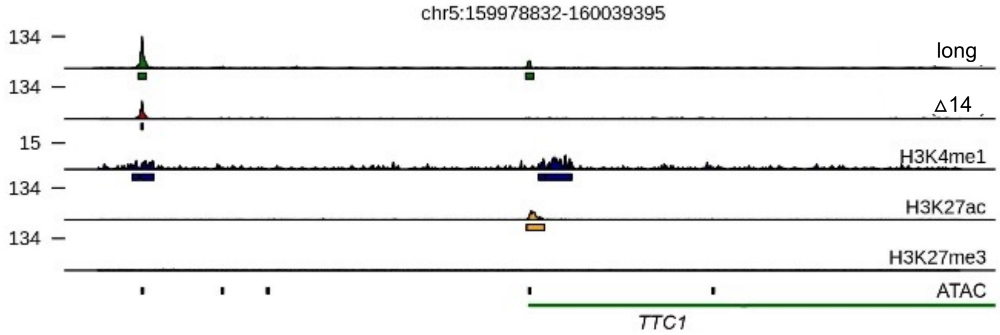

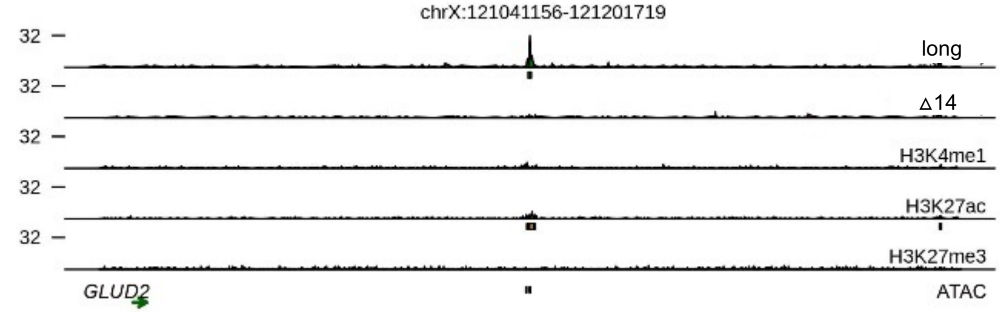
