## Supplementary material for "Alternative mRNA splicing controls the functions of the histone H3K27 demethylase UTX/KDM6A": Figure S5

| **location** | **mutation** | **Mutation type** | **amino acid** | **cancer** | **rate** |
|---|---|---|---|---|---|
| ***ICGC:*** | | | | | |
| **45060619** | CCT>CAT | missense | Pro>His | COAD | 1/420 |
| **45060619** | CCT>CTT | missense | Pro>Leu | UCEC | 1/531 |
| 45060677 | CAA>CAG | synonymous | Glu= | PACA | 1/391 |
| 45060728 * | AGT>AGC | synonymous | Ser= | UCEC | 1/351 |
| ***GnomAD***: | | | | | |
| 45060609 | GCA>ACA | missense /<br>splice region variant | Ala>Thr | - | 1/64391 |
